# Thermo-sensing and argonaute-dependent transcriptome remodelling in trypanosomes

**DOI:** 10.64898/2026.01.20.700697

**Authors:** Gustavo Bravo Ruiz, Michele Tinti, David Horn

## Abstract

Vector-borne parasites, such as African trypanosomes, experience temperature fluctuation due to fever symptomatic of infection, during developmental life cycle transitions, or due to diurnal shift. Mechanisms underpinning RNA-based thermo-sensing remain largely uncharacterised in these and other eukaryotic cells, however. Notably, trypanosomes exhibit almost exclusive polycistronic transcription, such that gene expression controls are dominated by post-transcriptional mechanisms mediated by mRNA-binding proteins and thousands of mRNA 3’-untranslated regions (3’-UTRs). Here, we quantify the transcriptomes and proteomes of bloodstream form *Trypanosoma brucei* following growth at 34°C, 37°C or 40°C for six hours. Approximately fifty genes encoding classical (co-)chaperones, with (UUA)_n_-rich 3’-UTRs, display the expected heat-shock response. The expression of approximately 1,000 additional transcripts is also correlated with temperature, and these transcripts have relatively long 3’-UTRs enriched in potentially complementary poly-purine tracts, poly-pyrimidine tracts, and palindromic sequences (P^5^-UTRs). To assess the potential impacts of mRNA secondary structure transcriptome-wide, we quantify mRNAs in cells lacking the central RNA interference nuclease argonaute (AGO1). Strains lacking AGO1 display increased retroposon expression, as expected, and strikingly abrogated thermo-regulation of transcripts with P^5^-UTRs. Thus, thermo-sensing involves argonaute-dependent transcriptome-remodelling in trypanosomes. We propose a post-transcriptional zipper hypothesis whereby access to regulatory motifs is controlled by temperature-sensitive mRNA secondary structure.

## INTRODUCTION

All cells and organisms modulate gene expression when exposed to variable temperatures. In the case of trypanosomatids, different hosts become infected during digenetic life cycles, such that these protozoal parasites typically encounter cooler or more variable temperatures in an insect vector and a typically higher, more constant, albeit also variable, temperature in a mammalian host. The African trypanosome, *Trypanosoma brucei*, for example, is transmitted by tsetse flies, and causes lethal and debilitating diseases, sleeping sickness in humans and nagana in livestock. Diurnal changes in temperature impact both insect and mammalian hosts; African cattle temperatures can vary by >2°C during the day, for example ^1^. In a human host, the haemolymphatic stage of trypanosomiasis is associated with recurrent pyrexia or fever ^2^, which can reach 41°C. Similarly affected vector-borne parasitic trypanosomatids include other African trypanosomes; the South American trypanosomes, *Trypanosoma cruzi*, which cause Chagas disease; and the *Leishmania* spp., which cause the leishmaniases ^3^.

The heat-shock response is broadly conserved among eukaryotes. Although it is characterised by reduced protein translation, expression of a specific set of (co-)chaperones increases; the classical ‘heat-shock proteins’ (HSPs) in particular ^4,5^. Proteins can become aggregated or misfolded at increased temperature, and these can be rescued or eliminated by cellular (re)folding or degradation activities ^6^. Trypanosomatids present a unique model system for exploring responses to a changing environment, since transcription is almost exclusively polycistronic in these Excavate protists, necessitating almost exclusive post-transcriptional gene expression control ^7^. Thus, increased transcription driven by heat-shock factors, as described in other eukaryotes ^8,9^, does not operate in trypanosomatids. Indeed, a *T. brucei* RNA binding protein, ZC3H11 ^10^, was found to stabilise target chaperone-encoding transcripts after heat-shock ^10^. More recently, a massive parallel reporter assay revealed many regulatory mRNA 3’-untranslated regions (3’-UTRs) in *T. brucei,* with adenine-rich poly-purine tracts linked to increased translation ^11^.

To mimic their natural environment, the developmentally distinct insect and mammalian bloodstream stages of *T. brucei* are typically grown in the laboratory at 27°C and 37°C, respectively. Prior transcriptomics analysis of insect stage *T. brucei* grown at 37°C or 41°C suggested a role for temperature in increasing expression of bloodstream stage upregulated transcripts ^12^ and our more recent *in silico* analyses suggested a potential role for uracil-rich poly-pyrimidine tracts in developmental controls ^11^. Pseudouridine modification on *T. brucei* spliceosomal small nuclear RNAs may also facilitate growth at elevated temperature ^13^. Prior proteomics analysis of bloodstream form *T. brucei* revealed key roles for phosphorylation of RNA binding proteins in co-ordinating the heat-shock response ^14^ and this analysis, which employed heat-shock at 41°C for one hour, revealed significantly increased abundance of HSP100 (ClpB1), but relatively few other changes in protein abundance. Proteomics analysis, following two-dimensional gel electrophoresis, has also been reported for *T. cruzi*, following growth at 28°C, 37°C or 42°C for three hours, revealing HSP70 and eighteen other proteins with increased intensity ^15^, while a more recent proteomics study on *Leishmania,* following growth at 25°C or 34°C, revealed several temperature-responsive HSPs ^16^.

Here, we use transcriptomics analysis, and proteomics analysis of bloodstream form *T. brucei* following growth at 34°C, 37°C or 40°C for six hours. We find that 3’-UTRs from post-transcriptionally regulated genes are enriched in low-complexity and potentially complementary sequences. Temperature-dependent increased expression of these transcripts is dependent upon the central RNA interference nuclease argonaute, indicating a role for RNA interference dependent regulation. The results reveal hundreds of thermo-sensitive transcripts in trypanosomes and suggest a zipper hypothesis for the regulation of both mRNA stability and for the progressive unmasking of regulatory sequences as temperature increases.

## RESULTS

### Transcriptomic and proteomic profiles of thermo-sensing in *T. brucei*

To identify temperature responsive transcripts and proteins in bloodstream form *T. brucei,* we cultured these cells at 34°C, 37°C or 40°C for six hours. Cell lysates and RNA samples were prepared and processed in triplicate for direct data-independent acquisition (directDIA) mass spectrometry and quantitative proteomic analysis, and for transcriptomic analysis, respectively (Fig. 1A, Supplementary Data 1). Heat-maps and principal component analyses revealed both temperature responsive and highly reproducible changes in protein and mRNA abundance (Supplementary Fig. 1A-C). Rather than displaying a common ‘stress’ response when temperature deviated from 37°C, changes in both mRNA and protein abundance correlated or inversely correlated with temperature (Fig. 1B). Scatterplots for the full set of 9,398 quantified transcripts and 5,974 quantified protein groups highlight the substantial and significant transcriptomic and proteomic responses to temperature shift from 37°C to 40°C (Fig. 1C). A comparison of the two datasets revealed substantial roles for both translation control and mRNA abundance control (Fig. 1D). Indeed, a cluster of genes registered significantly (FDR < 0.001) increased abundance for both mRNA and protein at 40°C and this more classical heat-shock response is discussed in more detail below.

**Fig 1:**
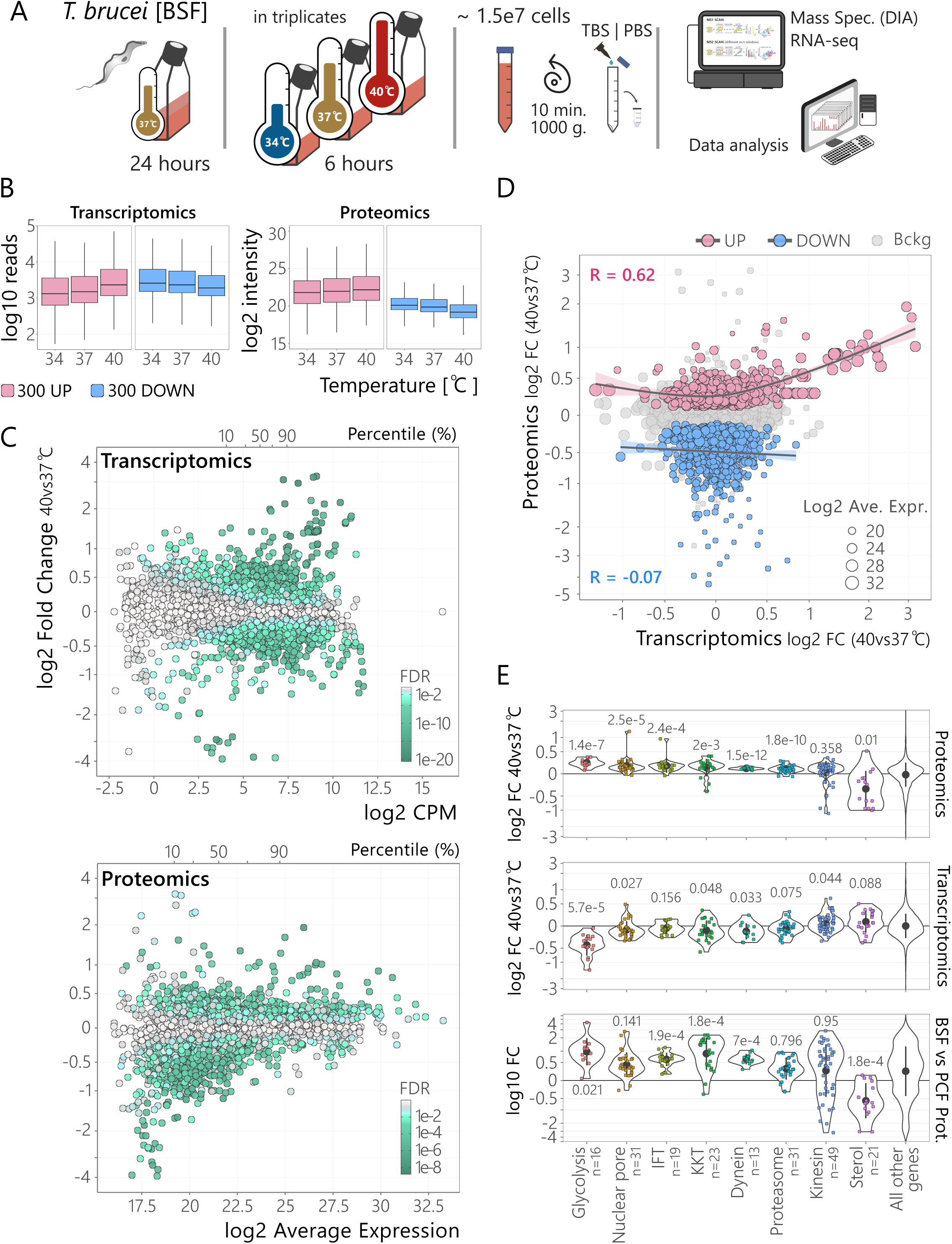
Proteomic and transcriptomic analysis of the *T. brucei* heat-shock response. **A,** The schematic shows sample preparation for proteomic and transcriptomic analysis. **B,** The boxplots show abundance for those top 300 mRNAs and proteins which report increased (red) or decreased (blue) abundance (FDR < 0.01) at 40°C compared to 37°C. **C,** The plots show mRNA (top panel; n = 9,398) and protein group (bottom panel; n = 5,974) abundance and fold changes at 40°C compared to 37°C. Log2 average expression or counts per million (CPM) reflects normalized mean intensity of replicates. The colour gradients represent the false discovery rate (FDR). **D,** The plot shows a comparison between the proteomic and transcriptomic data shown as log2 fold changes (FC) at 40°C compared to 37°C. Highlighted are proteins whose levels significantly increase (red) or decrease (blue) at 40°C (FDR < 0.01). For each subset, trend lines calculated using a Generalized Additive Model (GAM) and Pearson’s correlation (R) are indicated. **E,** The violin plots show fold-change at 40°C compared to 37°C for complexes and cohorts of proteins and transcripts (see Supplementary Data 1, sheet 5), and fold-change of protein abundance in the bloodstream form (BSF) relative to procyclic form (PCF) for the same groups [data from ^17^]. p-values, calculated using a *t*-test are shown for each group. IFT, intraflagellar transport protein; KKT, kinetoplastid kinetochore protein.

In terms of other likely primary impacts on cell physiology, abundance was significantly (FDR < 0.01) increased for 570 proteins and was significantly decreased for 742 proteins at 40°C. An analysis of protein cohorts and complexes indicated increased abundance at 40°C for glycolytic enzymes, nuclear pore complex components, intraflagellar transport proteins, kinetochore proteins, dyneins and most kinesins (Fig. 1E, upper panel). Perhaps surprisingly, these cohorts were typically moderately decreased in abundance at the mRNA level (Fig. 1E, middle panel). Notably, these same cohorts and complexes similarly displayed increased abundance in bloodstream form *T. brucei* relative to insect stage *T. brucei* (^17^, Fig. 1E, lower panel). These profiles, therefore, indicate developmental shifts in energy metabolism, and likely reflect the importance of maintaining intracellular traffic and macromolecular movement at increased temperature. Our observations also suggest that temperature plays a role in regulating the expression and abundance of these developmentally regulated proteins. Proteasomal proteins were also increased in abundance at 40°C, and in bloodstream form cells (Fig. 1E, upper and lower panels), likely reflecting increased protein turnover at increased temperature. We also note that typical markers of stress granules ^18^ were not increased in abundance at 40°C.

Thus, our identification of temperature responsive transcripts and proteins in bloodstream form *T. brucei* revealed substantial roles for both mRNA abundance control and translation control. Analysis of both datasets revealed a more classical heat-shock response (see below) and evidence that temperature makes a substantial contribution to developmental differences in relative protein abundance during the *T. brucei* life cycle.

### The classical heat-shock and chaperone response

Fifty-three genes registered significantly (FDR < 0.001) increased abundance at 40°C for both mRNA and protein (see Fig. 1D), and an analysis of these genes using Gene Ontology revealed significant enrichment of ‘protein folding’, ‘response to heat’ and ‘heat shock protein binding’ revealing a classical heat-shock response (Fig. 2A). Indeed, this cohort included many annotated heat-shock proteins (HSPs), and (co-)chaperones, which we describe in more detail here since very few have been experimentally assessed for temperature-dependent expression control in *T. brucei* previously. We first examined a core set of fifteen HSPs that are conserved in *T. brucei*, and the other parasitic trypanosomatids, *T. cruzi*, and *Leishmania major* (Fig. 2B; Supplementary Fig. 2; see Supplementary Data 1). All nine of those most abundant HSPs at 37°C were significantly (FDR < 0.002) increased in abundance following heat shock (Fig. 2C). Indeed, genes encoding the five most abundant HSPs (>27 log^2^ intensity at 37°C) are present in tandem arrays or multiple closely linked copies in the genome (see Supplementary Data 1); reflecting gene amplification as a mechanism to boost gene expression in trypanosomatids. Proteomics data for each heat-inducible HSP are shown in Figure 2D, revealing heat-inducible pairs of HSP90/70 proteins ^19^1 that localise to the cytosol, endoplasmic reticulum (ER), or mitochondrion; putative partner HSP110 ^20^ and localise to the cytosol or ER; and a mitochondrial HSP60. A robust ER response is notable here since prior analyses suggested the absence of an unfolded protein response in bloodstream form *T. brucei* ^21^. Thus, core sets of cytoplasmic, ER and mitochondrial HSPs display a robust heat-inducible transcriptomic and proteomic response in *T. brucei*.

**Fig 2:**
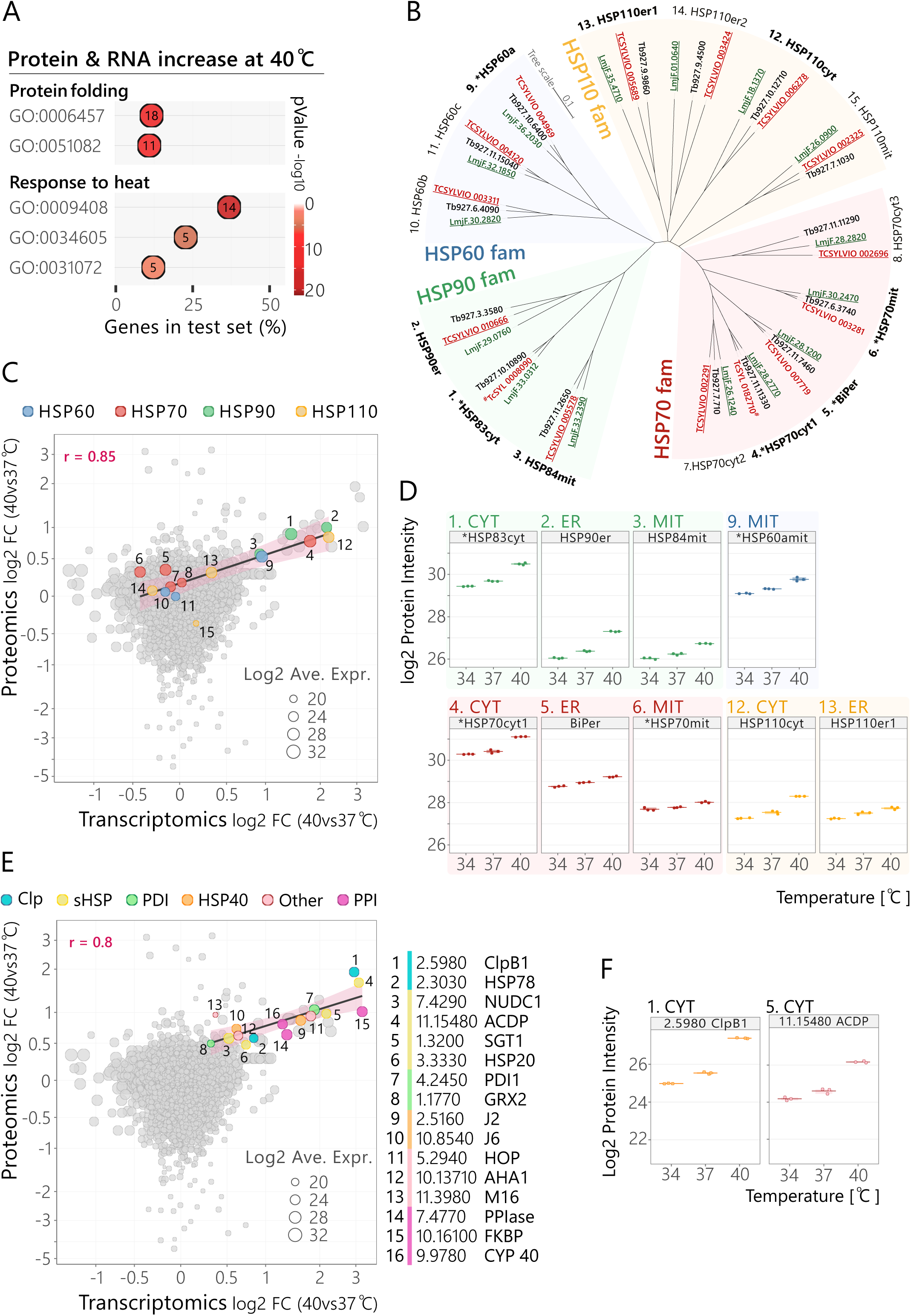
The classical heat-shock and chaperone response. **A,** Gene Ontology terms enriched among genes for which both protein and mRNA abundance was significantly increased (n = 53) at 40°C (FDR <0.001). The x-axis shows the frequency (percentage of the number of protein groups enriched in the analysis / total number associated with each GO term). Numbers within the circles indicate number of enriched genes associated with each GO-term; only considered if >4. **B,** The phylogenetic tree shows HSP60, HSP70, HSP90 and HSP110 family members from *T. brucei* (black), *T. cruzi* (red), and *Leishmania major* (green). Those nine HSPs that were significantly increased in abundance at 40°C (FDR <0.002) are highlighted in bold. An asterisk indicates proteins encoded by tandem copies in the *T. brucei* genome. Protein localization is indicated with; cyt, cytoplasmic; mit, mitochondrial; ER, endoplasmic reticulum. **C,** The plot shows a comparison between the proteomic and transcriptomic data as in Fig. 1D, highlighting the HSPs shown in Fig. 2B. A correlation line and the Pearson’s correlation value are shown. **D,** Replicate data for the heat-inducible HSPs showing relative abundance and highly reproducible changes in protein abundance in response to temperature change. The solid bar indicates the mean of the three replicates, and the box (where visible) indicates maximum and minimum values. Protein localization is indicated as in B. **E,** As in C but highlighting other heat-inducible (co-)chaperones, all with FDR <0.002. Clp-protease (Clp), protein disulfide-isomerase (PDI), small HSP (sHSP), peptidyl-prolyl isomerases (PPI), HSP40 family (HSP40-J2-J6), HSP organising protein (HOP), activator of HSP90 (AHA1), alpha-crystallin domain protein (ACDP), nuclear distribution C (NudC), suppressor of the G2 allele of SKP1 (SGT1), dithiol glutaredoxin 2 (GRX2), FK506 binding protein (FKBP), cyclophilin-type protein (CYP). **F,** Replicate protein data as in D and for ClpB1 and ACDP.

The remaining thirty-eight heat-inducible genes included several other small HSPs, isomerases, proteases, and co-chaperones (Fig. 2E-F). Two of these were HSP40-DnaJ family members, which comprise a large family ^22^, of >80 proteins in *T. brucei* and facilitate HSP70 action and substrate selection ^23^; Tbj2, previously shown to be heat-inducible, cytosolic, and essential ^24^, and Tbj6, thought to be involved in translation initiation in *T. cruzi* ^25^. Another two were among the ClpB/Hsp104 proteases of the HSP100 family, typically involved in dealing with aggregated and misfolded proteins ^6^; cytoplasmatic ClpB1 and mitochondrial HSP78. Mitochondrial HSP10, which works in conjunction with HSP60, as demonstrated in *Leishmania* ^26^, and a putative mitochondrial M16 metallopeptidase were heat-inducible, as were the cytosolic HSP organizing protein (HOP); an activator of HSP90 (AHA), which facilitates the transfer of HSP70 ‘clients’ to HSP90 ^27^; and three peptidyl-prolyl *cis-trans* isomerases (PPIs), including the cytosolic cyclophilin (CYP40) and FK506-binding protein (FKBP), which can accelerate protein (re)folding. Among nine α−crystallin domain (ACD) chaperones identified in trypanosomatids ^28^, which typically interact with partially folded proteins to prevent aggregation, four were heat-inducible; suppressor of G2 allele of SKP1 (SGT1), nuclear distribution C (NUDC1), HSP20; and ACDP; which appears to be trypanosomatid-specific. Finally, protein disulfide isomerase 1 (PDI1) ^29^ and the mitochondrial dithiol glutaredoxin (GRX2) ^30^ registered on the remaining list of thirty-eight heat-inducible genes.

### Temperature-dependent expression of promoter-adjacent transcripts

Before exploring temperature-dependent post-transcriptional gene expression controls in *T. brucei,* we assessed temperature-dependent impacts on polycistronic transcription units. *T. brucei* genes are organised almost exclusively in polycistronic units, which are approximately 120 kbp long on average and contain approximately fifty genes on average (see ^31^). Approximately 200 RNA polymerase II transcription start-sites display characteristics of highly dispersed or distributed promoters that lack conventional promoter-associated sequences ^32,33^. To investigate the impact of temperature on polycistronic transcription units, we plotted the transcriptome data relative to distance from transcription start-sites or transcription termination-sites (Fig. 3A). Transcripts from genes within 25 kbp of a transcription start-site displayed significantly increased abundance at 40°C relative to more distal genes (*p* = 4e^-111^) while transcripts from genes within 25 kbp of a transcription termination-site were not significantly different at 40°C (*p* = 0.4); notably, analysis of microarray data previously indicated increased expression of genes adjacent to transcription termination-sites, after 41°C heat-shock in insect-stage cells ^34^. We also note that protein abundance was not significantly impacted by distance from transcription start-sites (*p* = 0.07). Figure 3B shows how the abundance of transcripts adjacent to RNA polymerase II promoters was specifically increased in abundance at 40°C while the circular plot in Figure 3C shows significantly increased expression adjacent to the substantial majority of promoters genome-wide, with few examples of significantly reduced expression adjacent to these sites. Increased transcript abundance from promoter-adjacent genes in the absence of increased protein abundance may reflect increased transcription initiation and termination in these regions but failure to generate productive mRNA. Perhaps these regions, in association with transcriptional hubs ^35^, aberrantly assemble non-processive RNA polymerase II at elevated temperature. Nevertheless, we noted that most temperature-responsive genes were >25 kbp distal from promoters (see Fig. 3B-C), consistent with post-transcriptional control.

**Fig 3:**
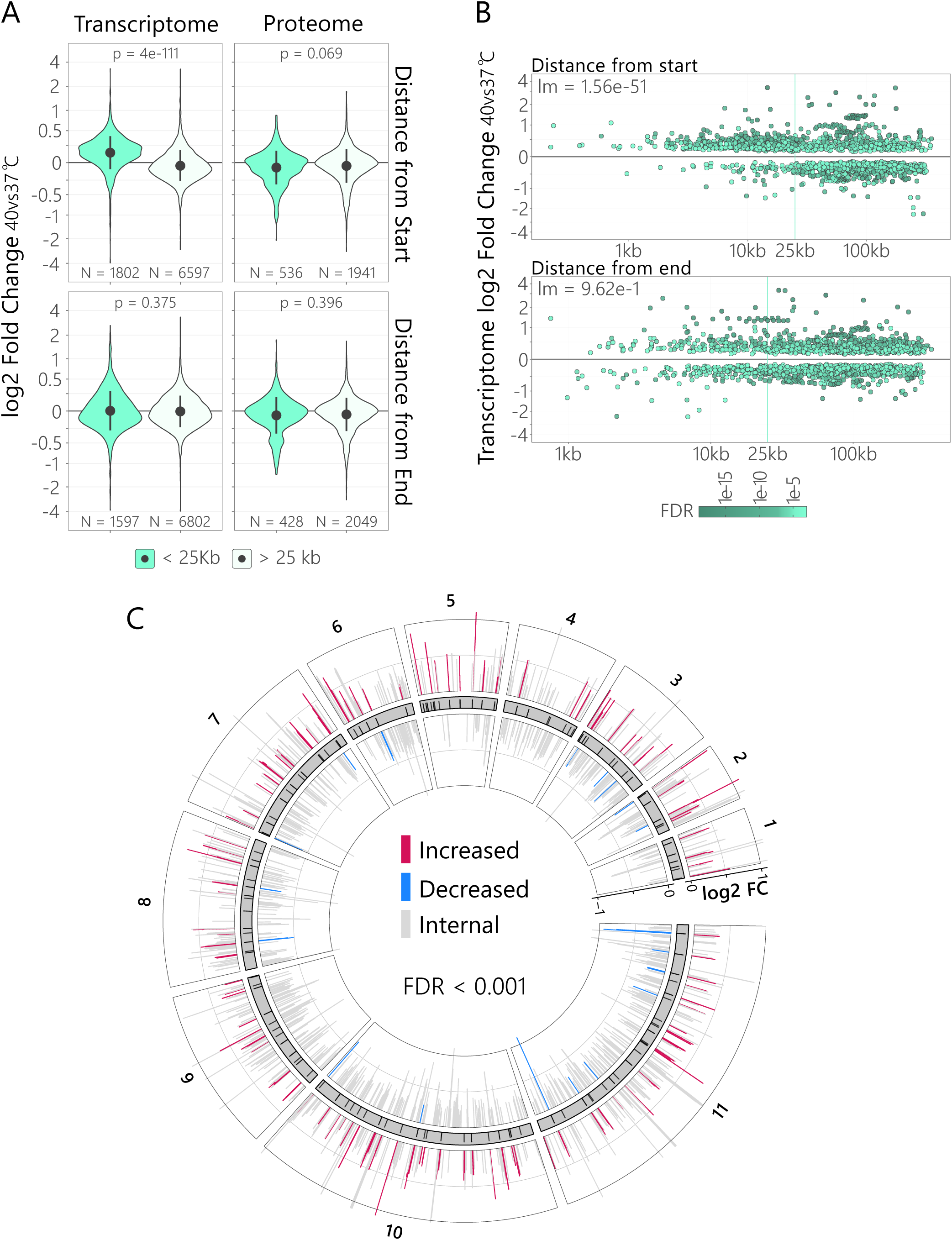
Temperature-dependent expression of promoter-adjacent transcripts. **A,** The violin plots show log2 fold changes in the transcriptomic and proteomic data at 40°C compared to 37°C, for genes < 25 Kb (green) or > 25 Kb (light green) from transcription start sites (top) or termination sites (bottom) in polycistrons. The circles indicate mean abundance, and vertical lines indicate SD. The p-values are from *t*-tests. **B,** The plots show log2 fold change at 40°C compared to 37°C for the transcriptome data relative to distance from transcription start sites (top) or termination sites start (bottom). Only significant changes (FDR < 0.001) are shown, and FDR is indicated by the colour gradient. The p-values are from linear regression (lm). **C,** The circular karyotype plot shows core chromosomes and log2 fold changes at 40°C compared to 37°C. Black bars show promoter positions, while coloured bars in the outer and inner circles show log2 fold changes at 40°C compared to 37°C for those transcripts that were < 10 Kb from a promoter and significantly different (FDR < 0.001); red = increased at 40°C, blue = decreased at 40°C. Other significantly different transcripts are shown in grey.

### Temperature responsive RNA-binding proteins

Before exploring mRNA 3’-untranslated regions (3’-UTRs) associated with temperature-dependent post-transcriptional gene expression controls, we assessed the full set of 231 mRNA-binding proteins (RBPs) in our transcriptomic and proteomic datasets (Fig. 4A). The most highly upregulated RBP at 40°C was ZC3H11, which in known to bind and upregulate mRNAs encoding heat-shock proteins ^10^. Other heat-inducible RBPs were RBP10, which is known to promote the bloodstream-form phenotype ^36^, and an uncharacterised RBP, Tb927.10.15610. In contrast, expression of RBP9 ^37^, and RBP17 was reduced at 40°C (Fig. 4A-B). These uncharacterised RBPs are new candidate regulators of the post-transcriptional response to temperature change.

**Fig 4:**
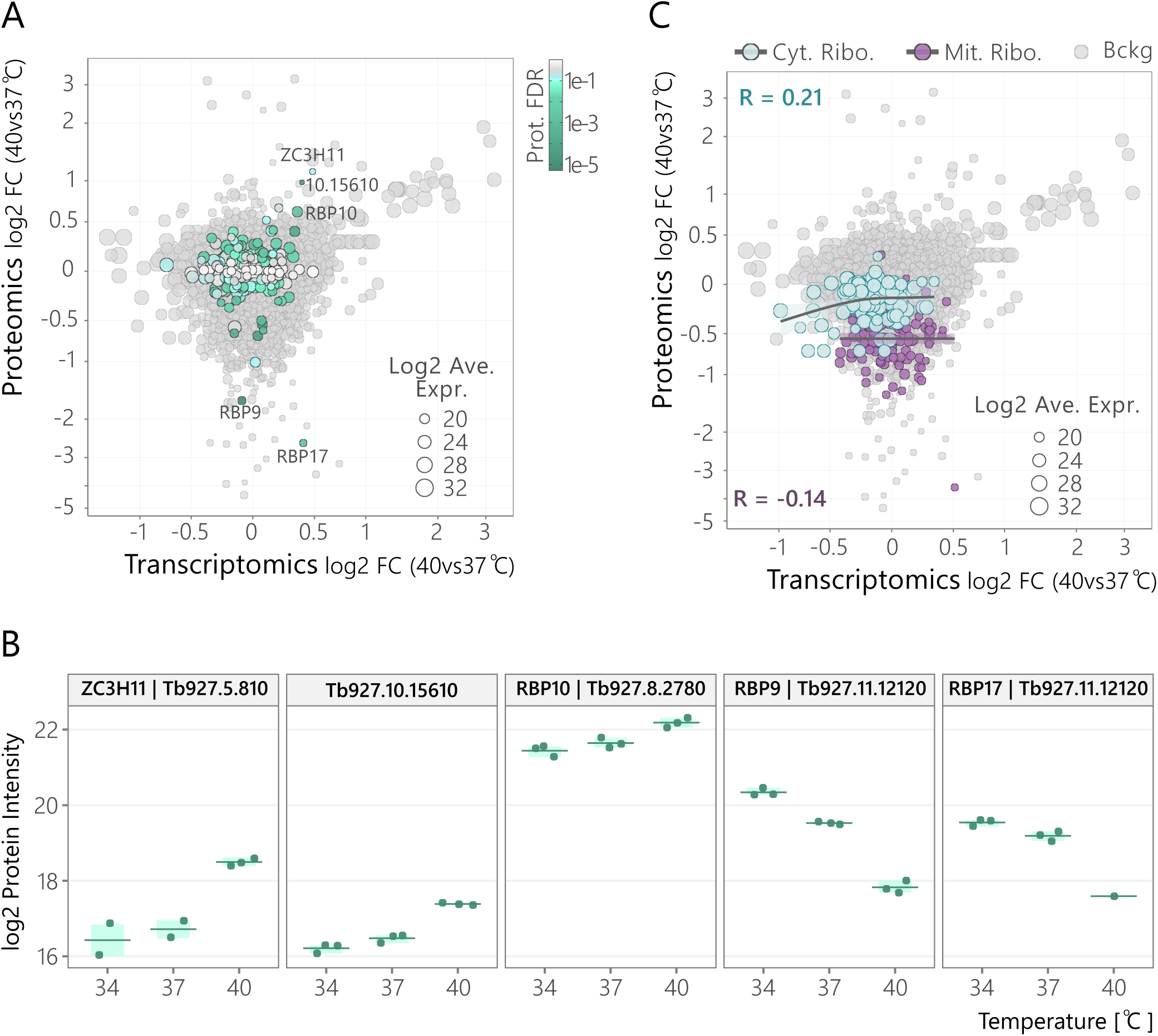
Temperature responsive RNA-binding proteins. **A,** The plot shows a comparison between the proteomic and transcriptomic data as in Fig. 1D, highlighting RNA binding proteins (RBPs; see Supplementary Data 1, sheet 5). FDR is indicated by the colour gradient. **B,** Replicate protein data as in Figure 2D for selected RBPs. **C,** The plot shows a comparison between the proteomic and transcriptomic data as in A, highlighting cytosolic ribosomal proteins (GO:0022626, cyan) and mitochondrial ribosomal proteins (GO:0005761, purple). For each subset, trend lines calculated using a Generalized Additive Model (GAM) and Pearson’s correlation (R) are indicated.

Since polysome abundance was previously reported to be decreased following heat-shock in *T. brucei* ^38^, we visualised both the mitochondrial and cytoplasmic ribosomal proteins in our datasets (Fig. 4C). The abundance of cytoplasmic ribosomal proteins was approx. 100-fold greater than the abundance of mitochondrial ribosomal proteins at all three temperatures analysed, but the abundance of both cohorts was reduced at 40°C. These results are consistent with the view that translation in both the mitochondrion and in the cytosol were reduced at 40°C. Indeed, those 377 proteins that were significantly (FDR < 0.001) reduced in abundance at 40°C (see Fig. 1C) were enriched for Gene Ontology terms associated with the ‘mitochondrion’, including ‘mitochondrial ribosomal subunits’, ‘kinetoplast’ (the structure containing the mitochondrial genome), ‘electron transport chain’ and ‘mitochondrial RNA processing’ (Supplementary Fig. 3).

### Thermo-sensitive transcripts are enriched for complementary motifs

Post-transcriptional expression controls in trypanosomatids are dominated by mRNA 3’-UTRs that selectively bind regulatory RBPs. To explore mRNA 3’-UTRs associated with temperature-dependent post-transcriptional gene expression controls in *T. brucei,* we assessed UTRs associated with thermo-sensitive genes in our transcriptomic and proteomic datasets. Specifically, we assessed the 3’-UTRs for the set of approximately fifty classical heat-shock genes, which registered a highly significant (FDR < 0.001) increase in both mRNA and protein abundance at 40°C, and for ninety-nine genes for which only protein abundance registered a highly significant (FDR < 0.001) increase at 40°C (Fig. 5A). We first assessed ^11^1 and found that while length for the classical heat-shock genes (average = 521 b) was not significantly different from a set of unregulated control transcripts (average = 507 b), the latter group of transcripts had significantly (*p* = 5.7e^-5^) longer 3’-UTRs (average = 996 b) (Fig. 5B).

**Fig 5:**
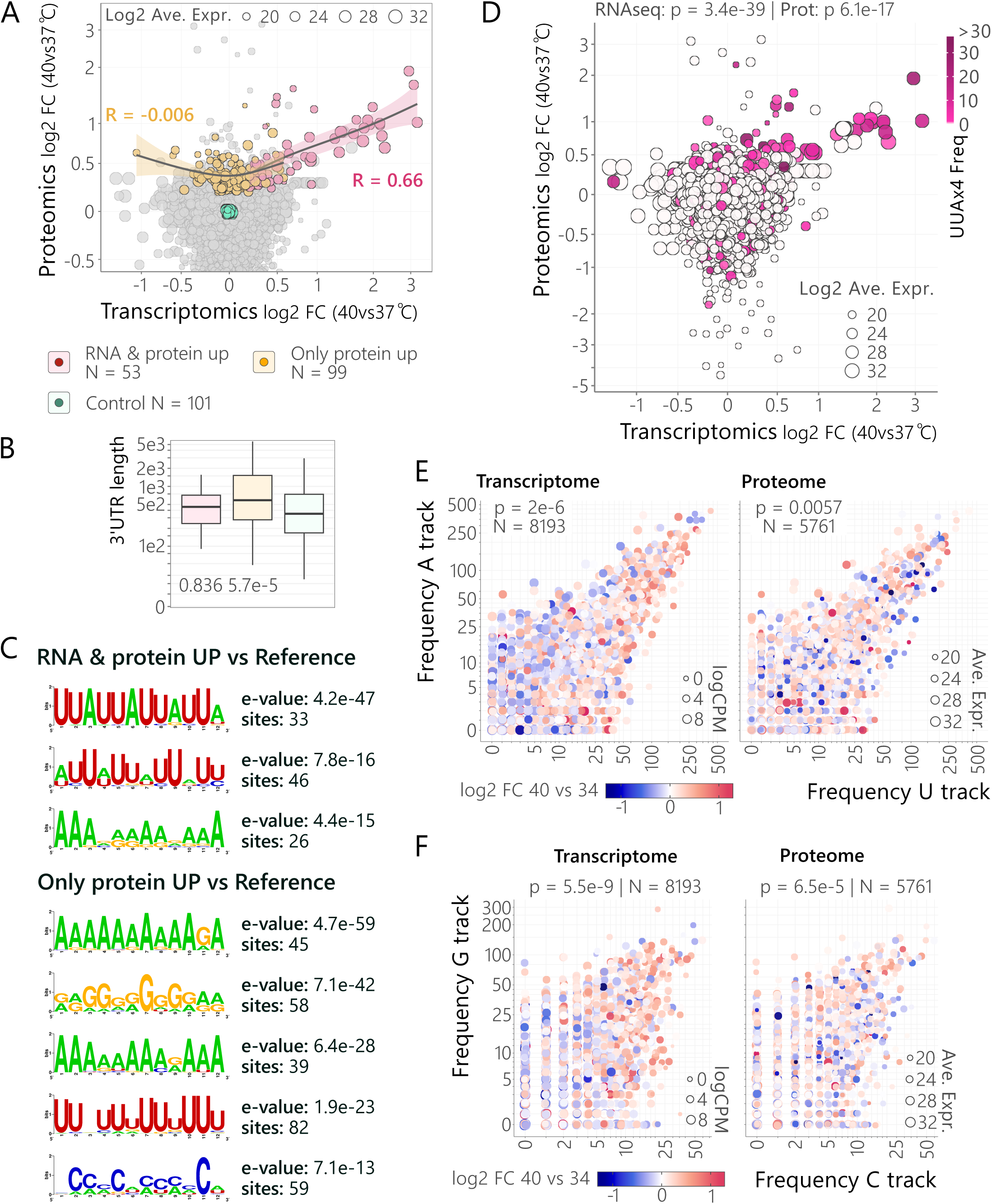
Thermo-sensitive mRNAs are enriched for complementary motifs. **A,** The plot shows a comparison between the proteomic and transcriptomic data as in Fig. 1D, highlighting genes for which both protein and mRNA abundance are significantly (FDR < 0.001) increased at 40°C (pink) or genes for which only protein abundance is significantly increased at 40°C (yellow). For each subset, trend lines calculated using a Generalized Additive Model (GAM) and Pearson’s correlation (R) are indicated. A reference set of temperature non-responsive genes is shown in green. **B,** The boxplot shows 3’-UTR length for the cohorts described in A. The p-values are from *t*-tests comparing each regulated cohort with the control set. **C,** MEME analyses revealed motifs enriched in the 3’-UTRs of the cohorts described in A relative to the reference set. Only motifs with e-value < 1e-10 are shown. **D,** The plot shows a comparison between the proteomic and transcriptomic data as in Fig. 1D. The colour gradient shows the frequency of (UUA)_4_ motifs (see C, top panel) in the 3’-UTRs. The p-values (p) are from linear regression analysis comparing the (UUA)_4_ frequency and transcriptomic (RNA-seq) or proteomic (Prot) log2 fold-change. **E,** The scatter-plots show comparisons between U-track and A-track frequency (see C, lower panel) in the 3’-UTRs. The colour gradient shows log2 fold change for transcripts (left) or proteins (right) at 40°C compared to 34°C. The p-values (p) are from multiple linear regression (MLR) analysis considering log2 fold change as the dependant variable and motif frequency as the independent variable. **F,** The scatter-plots show comparisons between C-track and G-track frequency (see C, lower panel) in the 3’-UTRs. Other details as in E.

We next assessed motif enrichment in the above 3’-UTRs and found that the classical heat-shock responsive transcripts were strongly enriched for a (UUA)_4_ motif (e = 4.2e^-47^) as well as an A-rich motif (Fig. 5C, upper panel). A similar ‘UUA’ motif was previously reported in the HSP70 3’-UTR, which is thought to be subject to regulation by the ZC3H11 RBP ^10^. These findings served to both validate our datasets and to substantially extend our understanding of the heat-shock response in *T. brucei*. We next turned our attention to the genes for which only protein abundance was significantly increased at 40°C. The longer than average 3’-UTRs in these transcripts revealed a distinct pattern of motif enrichment, with strong enrichment for polyPyrimidine-rich motifs (A-rich, e = 4.7e^-59^; G-rich, e = 7.1e^-42^) and potentially complementary polyPurine-rich motifs (U-rich, e = 1.9e^-23^; C-rich, e = 7.1e^-13^) (Fig. 5C, lower panel). Indeed, a second search for palindromic motifs revealed that these sequences were also specifically enriched in these 3’-UTRs (Supplementary Fig. 4); we refer to these 3’-UTRs as P^5^-UTRs since they are enriched for Poly-Purine, Poly-Pyrimidine, and Palindromic tracts.

The motifs detailed above are correlated with temperature-responsive expression for a relatively small numbers of genes, so we wondered whether these motifs could be predictive of temperature-responsive gene expression at a transcriptomic and/or proteomic scale. To address this question, we counted motifs in the full set of >8,000 annotated 3’-UTRs ^11^ and asked how these counts correlated with changes in expression at 40°C. The (UUA)_4_ motif was indeed predictive of both increased mRNA (*p* = 3.4e^-39^) and protein (*p* = 6.1e^-17^) abundance at a transcriptomic and proteomic scale (Fig. 5D). We next assessed how frequency of the potentially complementary A-rich and U-rich motifs correlated with changes in expression at 40°C. The frequency of these motifs was indeed significantly predictive of temperature-dependent responses for both the transcriptome (*p* = 2e^-6^) and the proteome (*p* = 5.7e^-3^) (Fig. 5E). Indeed, this was also true for potentially complementary G-rich and C-rich motifs, and also for both the transcriptome (*p* = 5.5e^-9^) and the proteome (*p* = 6.5e^-5^) (Fig. 5F). These results suggested a model whereby complementary sequences within P^5^-UTRs form double-stranded RNA (dsRNA) that is modulated by temperature and that impacts access to other regulatory sequences.

### AGO1-dependent thermo-sensing and transcriptome-remodelling

Our results above suggested a model whereby P^5^-UTRs form dsRNA that impacts gene expression in a temperature-dependent manner. RNA interference operates in *T. brucei* and depends upon the central small interfering RNA-programmed mRNA nuclease, argonaute 1 (AGO1) or slicer ^39^. We reasoned that dsRNA formed by P^5^-UTRs could be targeted by the RNAi machinery and impact thermo-sensing. To test this hypothesis, we used Cas9-based editing to knockout *AGO1* in bloodstream form *T. brucei*. Specific knockout was confirmed in two independent *ago1* null strains using whole genome sequencing (Fig. 6A). We then cultured both wild-type and *ago1* null strains at 34°C, 37°C or 40°C for six hours and generated transcriptome data for each sample (Supplementary Data 1). Since AGO1 is known to control retroposon transcripts in *T. brucei* ^39^, we first assessed impacts on these transcripts. As expected, several hundred INGI retroposon (*p* = 1.3e^-17^) and reverse transcriptase genes (*p* = 6e^-8^) reported significantly increased transcript abundance in the *ago1* null strains at 37°C (Fig. 6B), and indeed at all three temperatures tested; RIME (Ribosomal Interspersed Mobile Element) and RHS (Retrotransposon Hot Spot) transcripts ^40^ failed to report significantly increased expression in the *ago1* null strains, however.

**Fig 6:**
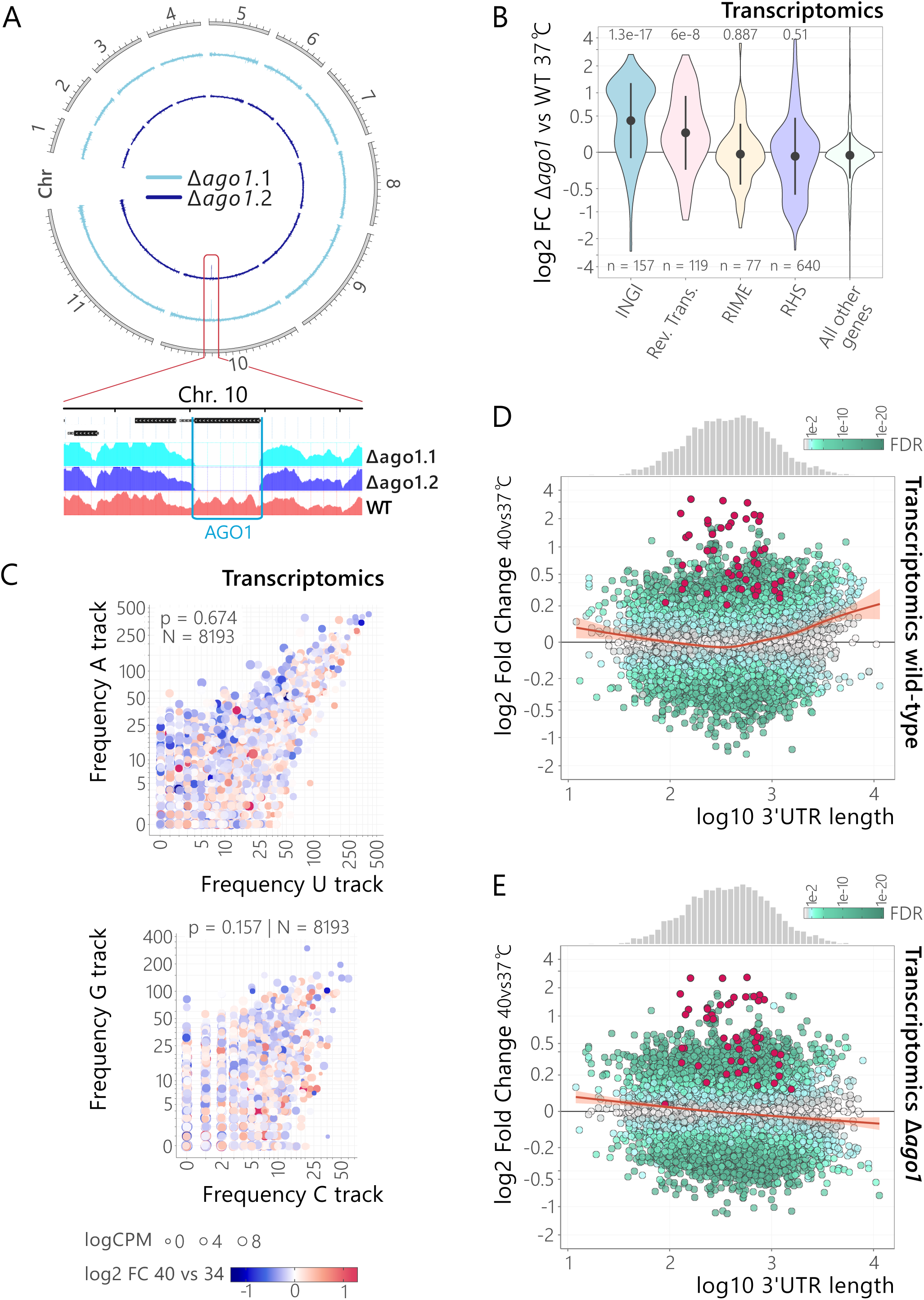
AGO1-dependent thermo-sensing and transcriptome-remodelling. **A,** The circular plot shows whole genome sequencing data for two independent *ago1* null strains compared to a wild-type control strain, revealing specific knockout of the *AGO1* gene on chromosome 10. The lower, zoomed panel shows precise *AGO1* deletion. **B,** The violin plot indicates increased retroposon transcript expression in *ago1* null cells at 37°C; INGI-containing transcripts and reverse transcriptases (Rev. Trans.). RIME-containing transcripts, and retrotransposon hot spot (RHS) are also shown (see Supplementary Data 1, sheet 5). The circles indicate mean abundance, and vertical lines indicate SD. The p-values are from *t*-tests relative to all other genes. **C,** The scatter-plots show comparisons between U-track and A-track frequency (upper panel) or C-track and G-track frequency (lower panel) in 3’-UTRs. The colour gradient shows log2 fold change for transcripts at 40°C compared to 34°C in *ago1* null cells. Other details as in Fig. 5E. **D,** The scatter-plot shows log2 fold-change for transcripts at 40°C relative to 37°C in relation to 3’-UTR length in wild-type cells (see dataset in Fig. 1C). The colour gradient represents false discovery rate (FDR). The trend line was calculated using a Generalized Additive Model (GAM). Highlighted in red are the heat-shock genes for which both protein and mRNA abundance were significantly (FDR < 0.001) increased at 40°C (pink in Fig. 5A). The upper histogram shows 3’-UTR length distribution across the dataset (50 bins). **E,** The scatter-plot shows log2 fold-change for transcripts at 40°C relative to 37°C in relation to 3’-UTR length in *ago1* null cells. Other details as in D.

We next turned our attention to P^5^-UTRs and, as above, assessed how frequency of potentially complementary motifs correlated with changes in expression at 40°C. The correlation between these motifs and the temperature-dependent transcriptome response observed above (Fig. 5E) was strikingly abrogated in *ago1* null cells; indeed, this was true for potentially complementary A-rich and U-rich motifs, and for G-rich and C-rich motifs in 3’-UTRs (Fig. 6C). Thus, we observe argonaute-dependent thermo-regulation of transcripts with P^5^-UTRs. To further visualise the impact of thermo-regulation at the transcriptomic scale, and the role of AGO1, we plotted 3’-UTR length against transcript fold-change at 40°C v 37°C. These plots reveal substantially increased abundance of approximately 1,000 transcripts with relatively long 3’-UTRs at 40°C in wild-type *T. brucei* (Fig. 6D) and striking abrogation of this thermo-sensing response in *ago1* null cells (Fig. 6E). Notably, the classical heat-shock response remained intact in *ago1* null cells (Fig. 6D-E), consistent with the view that ZC3H11 controls this response by binding (UUA)_4_ elements in these 3’-UTRs.

We next wondered whether 5’-UTRs might also contribute to thermo-sensing, and used a machine learning approach to determine whether features within 5’-UTRs, 3’-UTRs or both combined, correlated with observed transcriptomic or proteomic changes between 40°C and 34°C. The models indicated a substantial contribution of 3’-UTRs both transcriptome control and proteome control, as expected (see Fig. 5E-F), and as indicated by Area Under the Curve (AUC) values of 0.78 and 0.67, respectively (Fig. 7A). The models also indicated a substantial contribution of 5’-UTRs, but only for the proteomic dataset in this case, suggesting that 5’-UTRs selectively impact translation (Fig. 7A). Indeed, 5’-UTR length for genes for which only protein abundance was significantly increased at 40°C had significantly (*p* = 0.023) longer 5’-UTRs (average = 225 b) relative to the set of unregulated control transcripts (average = 136 b), while 5’-UTR length for the classical heat-shock genes (average = 120 b) was not significantly different (Fig. 7B).

**Fig 7:**
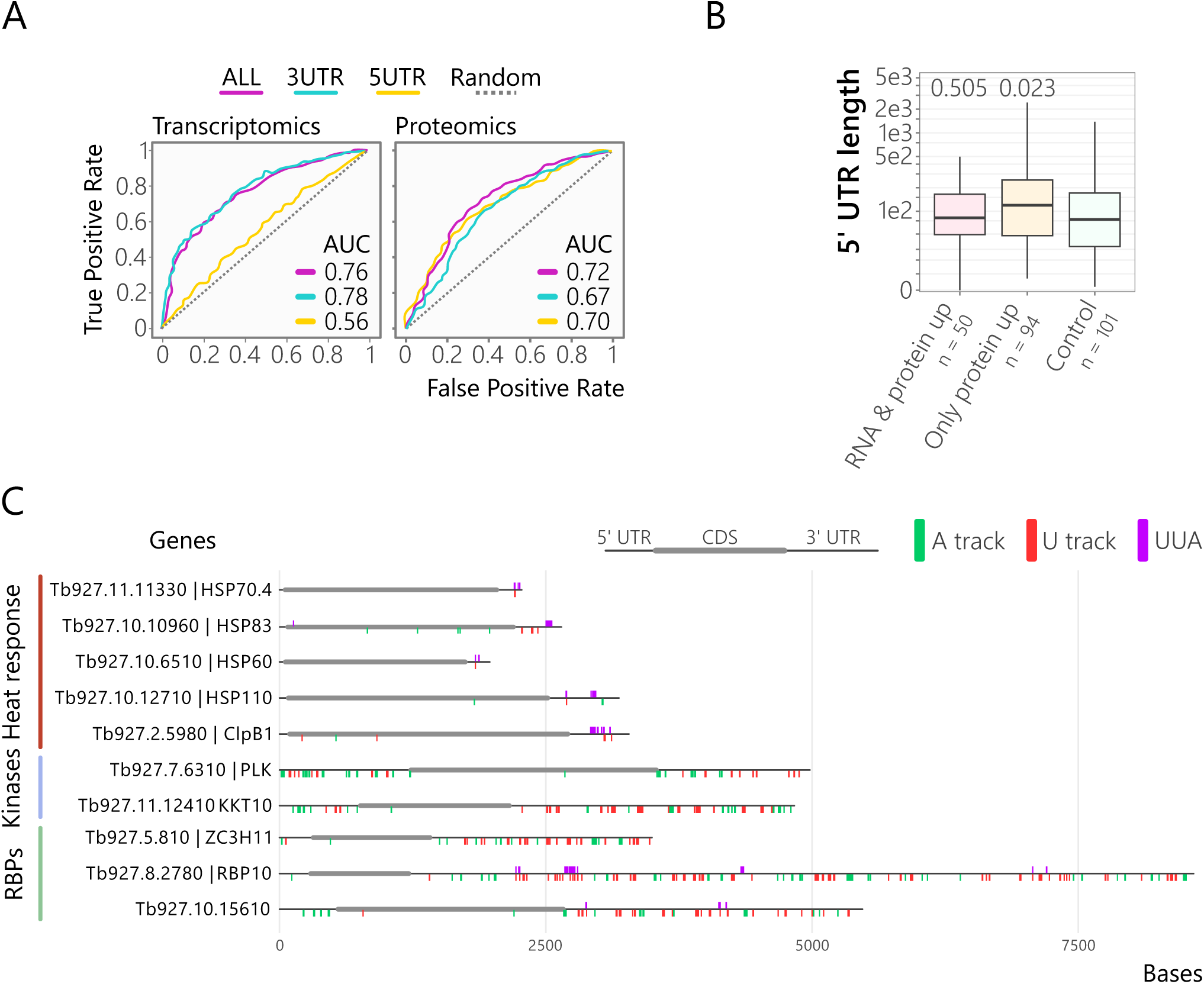
Machine learning analysis suggests a role for 5’-UTRs in translation control. **A**, Machine learning analysis. The receiver operating characteristic (ROC) curves show the predictive performance of UTR embedding approaches for distinguishing ∼300 up- or down-regulated genes following heat shock treatment using proteomic (left) or transcriptomic (right) data. Performance is evaluated using three UTR embedding strategies: 5’-UTR only (5UTR, orange), 3’-UTR only (3UTR, cyan), and combined 5’- and 3’-UTRs (ALL, magenta). Area under the curve (AUC) values are indicated in parentheses. Dashed grey line represents random classifier performance (AUC = 0.5). **B,** The boxplot shows 5’-UTR length for the cohorts described in Fig. 5B. The p-values are from *t*-tests comparing each regulated cohort with the control set. **C,** Motif location along the mRNA transcripts for selected exemplar genes. The A-track, U-track and (UUA)_4_ motifs are shown and the exemplars include classical heat-shock genes, and heat-inducible kinases and RBPs.

Schematic maps for a set of exemplar transcripts serve to illustrate the locations of (UUA)_4_ and/or potentially complementary A-rich and U-rich motifs described above (Fig 7C). The (UUA)_4_ motif is selectively enriched in the 3’-UTRs of *HSP* transcripts (*HSP20, HSP60, HSP70, HSP83, HSP110 and ClpB1* are shown), as can also be seen in Figure 5D, while A-rich and U-rich sequences are present in both 5’-UTRs and 3’-UTRs; two kinases (*KKT10* and *PLK1*) and three RBPs (*ZC3H11, RBP10* and *10.15610*) are shown. The KKT10/CLK1 kinetochore kinase and the polo-like kinase, display an abundance of A-rich and U-rich sequences in both their 5’- and 3’-UTR and differential expression of these and other potential regulators could provide an explanation for the mitosis and chromosome segregation defects described in *ago1* null insect stage *T. brucei* ^41^; although life-cycle progression was reported by others to be normal in *ago1* null cells ^42^. We note that both (UUA)_4_ and A-rich and U-rich sequences are present in the 3’-UTR of *RBP10,* which is known to promote the bloodstream-form phenotype ^36^ while only A-rich and U-rich sequences are present in the 3’-UTR of *ZC3H11;* suggesting the thermo-sensitive RBP10 mRNA may be subject to control by both ZC3H11 and AGO1. We conclude that, in addition to a more classical heat-shock response, both longer 5’-UTRs and 3’-UTRs (specifically P^5^-UTRs) can contribute to mRNA secondary structure and AGO1-dependent thermo-sensing in *T. brucei*.

## DISCUSSION

Post-transcriptional gene expression controls in trypanosomatids are dominated by mRNA 3’-untranslated regions that bind regulatory RNA-binding proteins. We recently reported a massive parallel reporter assay which revealed thousands of regulatory 3’-untranslated regions in *T. brucei,* linking adenine-rich poly-purine tracts to increased translation ^11^. *In silico* analyses reported in that study, also suggested a role for uracil-rich poly-pyrimidine tracts in developmental controls, potentially driven by temperature fluctuation and we have now used transcriptomics and proteomics analyses to explore the heat-shock and thermo-sensing response in the African trypanosome. Our multi-omics analysis revealed a classical heat-shock response involving substantial roles for both mRNA abundance control and translation control. The analysis also supported the view that temperature makes a substantial contribution to developmental differences in relative protein abundance during the *T. brucei* life cycle. Transcripts derived from promoter-adjacent regions are also elevated at increased temperature indicating impact at polycistronic transcription start-sites. Most notably, we find increased expression of transcripts with long, low-complexity 3’-untranslated regions that is dependent on the central RNAi nuclease, AGO1. We suggest a zipper hypothesis for transcriptome remodelling, whereby temperature-dependent secondary structure impacts AGO1-dependent regulation.

Although trypanosomatid heat-shock proteins are potential drug targets ^43,44^, or diagnostic markers ^45^, that impact differentiation ^46^, and drug sensitivity ^47,48^, the heat-shock response in African trypanosomes had not previously been described in detail at the transcriptomic or proteomic level. HSPs play important roles in protein folding, secretion, disaggregation, and degradation control. Expression of these proteins is increased in response to elevated temperature and other stresses, and we now describe a robust heat-shock response in *T. brucei*. We assessed a set of HSP60, 70, 90 and 110 chaperones that are conserved among the trypanosomatids, and find evidence for an unfolded protein response, previously thought to be absent in bloodstream form *T. brucei* ^21^. Several mitochondrial chaperones and co-chaperones are heat-induced, which is notable given simultaneously reduced expression of many other mitochondrial proteins. Indeed, both mitochondrial and cytoplasmic ribosomal protein abundance is reduced by heat-shock, consistent with globally reduced translation. Proteins involved in glycolysis, and cellular traffic and dynamics are not subject to this general trend, however. On the contrary, proteins associated with these functions display increased expression following heat-shock.

Unusually, gene expression control in trypanosomatids operates almost exclusively post-transcription, and consistent with this, we identify heat-responsive RNA-binding proteins known to execute post-transcriptional controls; most notably, ZC3H11, a known regulator of the heat-shock response ^10^, as well as other new candidate regulators of the post-transcriptional response to temperature change.

A particular focus in this study was to determine whether *T. brucei* 3’-untranslated regions play a role in thermo-sensing. We recently linked adenine-rich poly-purine tracts to increased translation, while *in silico* analyses suggested potential interactions with uracil-rich poly-pyrimidine tracts in developmentally regulated transcripts ^11^. Similar, RNA-based thermo-sensors may be more pervasive than previously appreciated, and have been described in bacteria ^49^ and plants ^50^, while the transcriptomes of rice ^51^ and malaria parasites, *Plasmodium falciparum* ^52^ are also thought to be subject to thermoregulation. More specifically, it has been suggested that translation regulatory elements in the 3’-untranslated region of the *Leishmania* Hsp83 transcript are more accessible at elevated temperature ^53^. Our current study shows that long P^5^-untranslated regions, enriched for Poly-Purine, Poly-Pyrimidine, and Palindromic tracts are thermo-sensitive and are increased in relative abundance at 40°C. We further demonstrate that this response does not operate in the absence of the major RNA interference nuclease, argonaute, or AGO1.

RNA interference in *T. brucei* is triggered by long dsRNA, which is cleaved by dicer to yield 20–30 nt small interfering RNAs (siRNAs), which then program AGO1, or slicer, to target complementary transcripts ^54^. Notably, *T. brucei* cells express both cytoplasmic and nuclear dicer-like proteins ^55^ and we observed a significant (1.5e^-5^; see Supplementary Data 1) reduction in expression of the nuclear dicer-like protein (DCL2) at 40°C. Thus, although thermo-sensitive transcriptome-remodelling described here is AGO1-dependent, multiple other trypanosomatid RNA nucleases, including DCL2, may interact with P^5^-UTRs and other RNA secondary structures.

We conclude that *T. brucei* cells substantially remodel their transcriptome and proteome in response to heat-stress. We observe argonaute-dependent transcriptome-remodelling, consistent with thermo-sensitive annealing of complementary sequences in P^5^-UTRs. Our findings support a zipper hypothesis, whereby temperature-sensitive mRNA secondary structure controls access to regulatory motifs that underpin post-transcriptional gene expression controls.

## METHODS

### *T. brucei* growth and manipulation

Bloodstream form Lister 427 *T. brucei* and derivatives, including 2T1^T7-Cas9^ ^56^, were cultured in HMI-11 (Gibco) supplemented with 10% fetal bovine serum (Sigma) and with 5% CO_2_ in a humidified incubator, at 37°C unless stated otherwise. Transfections were performed using 2.5 x 10^7^ cells, resuspended in homemade transfection buffer ^57^ with appropriate linearised plasmid DNA and electroporated using a Nucleofector system (Lonza) set on program Z-001 as described ^58^. After 4 to 6 h, transformants were cloned by limiting dilution and selected with the appropriate antibiotic(s).

### Transcriptomic analysis

*T. brucei* extracts from wild-type (Lister 427) and Δ*ago1*::NPT (two clones, see below) were obtained after growth of ∼ 1.5 x 10^7^ cells for 6 h at 34°C, 37°C or 40°C in triplicate. Cultures were centrifuged for 10 min at 1000 g and the cells were washed twice with PBS. RNA extraction was performed using the RNAeasy Mini Kit (Qiagen) following the manufacturer’s instructions. After RNA quantification by BioDrop (Fisher Scientific), 2-3 μg were sent for RNA-seq analysis on a DNBSEQ-G400 platform (BGI, Hong-Kong) where raw sequencing data was processed through a standard pipeline. Initial quality control of fastq files was performed using FastQC, followed by adapter trimming and quality filtering with Fastp (0.20.0) ^59^. Processed reads were aligned to the reference TREU927 genome or Lister 427_2018 genome (TriTrypDB v68) ^60^ using Bowtie2 (2.3.5) ^61^ with ’--very-sensitive-local’ parameters. The resulting alignments were processed with SAMtools (1.9) ^62^ for sorting and indexing, and PCR duplicates were marked using Picard MarkDuplicates (2.22.3) ^63^. Read counts per coding sequence were quantified using featureCounts (1.6.4) ^64^ with parameters accounting for multi-mapping reads (-M) and overlapping features (-O) and configured to count only reads where both ends were mapped (-B) and to exclude chimeric reads that mapped to different chromosomes (-C). The datasets were then used as input for differential expression analysis using the edgeR package (3.28) ^65^ in R (3.6.1). We retained only genes that had a minimum count of 10 reads in at least one sample and a minimum total count of 30 reads across all samples. The cqn package (1.32) ^66^ was used to compute gene length and GC content bias. The resulting offsets were incorporated into the DGEList object before computing dispersions. We fitted our data to a generalized linear model (GLM) using the quasi-likelihood (QL) method via glmQLFit and performed differential expression testing using the quasi-likelihood F-test through the glmQLFTest function. The output from this statistical test was then processed using the topTags function, which extracted the complete set of test results. We configured topTags to return all tested genes (n=Inf) without applying any sorting (sort.by=“none”), ensuring that the original gene order was preserved in the output. For multiple testing correction, we employed the Benjamini-Hochberg (BH) procedure (adjust.method=“BH”) to control the false discovery rate.

### Mass spectrometry analysis

To perform proteomic assay under heat stress conditions, *T. brucei* cell extracts were prepared for mass spectrometry. Cells were obtained after growth of ∼ 1.5 x 10^7^ cells for 6 h at 34°C, 37°C or 40°C in triplicate. The cells were centrifuged for 10 min at 1000 g, washed with PBS and resuspended in 100 μL of a solution containing 5% SDS and 100 mM triethylammonium bicarbonate. The samples were submitted to the Fingerprints Proteomics Facility at the University of Dundee and processed using trypsin, μBCA (bicinchoninic acid), strap processed, quality controlled, and peptide quantified. 200 μg from each sample was used, and the final peptide quantification yielded between 72 and 120 μg. For liquid chromatography-mass spectrometry (LC-MS) analysis, 1.5 μg of each sample was injected onto a nanoscale C_18_ reverse-phase chromatography system (UltiMate 3000 RSLC [rapid-separation liquid chromatography] nano; Thermo Scientific) and then electrosprayed into a Q Exactive HF-X mass spectrometer (Thermo Scientific). For liquid chromatography, buffers were as follows: buffer A was 0.1% (vol/vol) formic acid in MilliQ water, and buffer B was 80% (vol/vol) acetonitrile and 0.1% (vol/vol) formic acid in MilliQ water. Samples were loaded at 10 μL/min onto a trap column (100-μm by 2-cm PepMap nanoViper C_18_ column, 5 μm, 100 Å; Thermo Scientific) equilibrated in 0.1% trifluoroacetic acid (TFA). The trap column was washed for 3 min at the same flow rate with 0.1% TFA and then switched in line with a Thermo Scientific resolving C_18_ column (75-μm by 50-cm PepMap RSLC C_18_ column, 2 μm, 100 Å). The peptides were eluted from the column at a constant flow rate of 300 nL/min with a linear gradient from 3% buffer B to 6% buffer B for 5 min and then from 6% buffer B to 35% buffer B for 115 min and, finally, to 80% buffer B for 7 min. The column was then washed with 80% buffer B for 4 min and re-equilibrated in 35% buffer B for 5 min. Two blanks were run between each sample to reduce carryover. The column was kept at a constant temperature of 40°C. The data were acquired using an easy-spray source operated in positive mode with spray voltage at 2,500 kV and the ion transfer tube temperature at 250°C. The MS system was operated in DIA mode. A scan cycle comprised a full MS scan (m/z range from 350 to 1,650), with the RF lens at 40%, the automatic gain control (AGC) target set to custom, the normalized AGC target set at 300, the maximum injection time mode set to custom, the maximum injection time at 20 ms, and source fragmentation disabled. The MS survey scan was followed by tandem MS (MS/MS) DIA scan events using the following parameters: multiplex ions set to false; collision energy mode set to stepped; collision energy type set to normalized; high-energy collisional dissociation (HCD) collision energies set to 25.5, 27, and 30; orbitrap resolution at 30,000; first mass at 200; radio frequency lens at 40; AGC target set to custom; normalized AGC target at 3,000; and maximum injection time of 55 ms. Data for both MS and MS/MS scans were acquired in profile mode. Mass accuracy was checked before the start of sample analysis.

### Proteomic data analysis

We generated a spectral library based on the predicted protein sequences for *T. brucei* TREU927 or Lister 427 2018 strains, sourced from TriTrypDB (version 51) ^60^. DIA-NN (version 1.8.1) ^67^ was employed to process the raw mass spectrometry data. The data were analysed using default settings, leveraging both Unrelated Runs and Match Between Runs (MBR) functionalities. After DIA-NN processing, the output was transformed into a Protein Groups quantification table using the DIA-NN ^67^ R package, adhering to a 0.01 False Discovery Rate. Protein groups identified based on a single peptide or exhibiting zero intensity were designated as NaN (Not a Number) for quantification. Any group where all quantifications were NaN was omitted from further analysis. To normalize the data, column values underwent a log_2_ transformation post-median equalization. For conditions where all values were missing, they were interpreted as missing-not-at-random and replaced with minor random values approximating the detection threshold. In contrast, other missing values were deemed missing-at-random and substituted using an iterative imputer from the scikit-learn Python library ^68^. Finally, a Principal Component Analysis (PCA) was executed using the same scikit-learn library using two dimensions. Differential quantification analysis was conducted using the limma package in R ^69^. A linear model was fitted using the lmFit() function, followed by Empirical Bayes Statistics for Differential Expression computed with the ebayes() function. Adjusted *p*-values were calculated using the topTable() function with the Benjamini & Hochberg (BH) correction method. Analysis of variance (ANOVA) was carried out using the oneway.anova function from the HybridMTest package in R ^70^. The p.adjust function was employed to calculate adjusted *p*-values, again using the Benjamini & Hochberg (BH) correction.

### Generation of *ago1* null strain

The *AGO1* (Tb927.10.10850) gene was replaced by the neomycin-phosphotransferase (*NPT*) marker conferring resistance to G418 and generating the Δ*ago1*::NPT strain. As parental strain, we used the 2T1^T7-Cas9^ as described ^56^. First, a T7Cas9^AGO1_sgRNA^ strain was generated by introduction of specific AGO1 sgRNA designed manually. The sgRNA oligonucleotide pairs used were AGO1_gRNA-F (agggGGGGAGGGTCGTGAAGTTCT) and AGO1_gRNA-R (aaacAGAACTTCACGACCCTCCCC). The Δ*ago1*::NPT strains were generated by introducing specific targeted DNA templates (∼ 8 μg) to each T7Cas9^AGO1_sgRNA^ strain, each induced with tetracycline (1 μg/ml) for 3 h to activate Cas9 expression. After transfection, cells were grown in the presence of G418 (2.5 μg/ml) and tetracycline (1 μg/ml). Specific targeting DNA templates were obtained by PCR, using a plasmid containing the *NPT* cassette, and the primer pairs NPT_5UTR-AGO1 (catcaactaaaaaagcaaacaagcaacctaataaaaaggatttaaataaaATGATTGAACAAGATGGATTGC AC) and NPT_3UTR-AGO1 (tcacacacattgcaccttcccaaagttgctttccccggagaagcggtggtGATAAGCTTGATGGCCACGATG). The primers incorporate 50 bp corresponding to the 5’- and 3’-UTR sequence of each target locus, to promote homologous recombination.

### Whole genome sequencing

Genomic DNA from two Δ*ago1*::NPT strains and the 2T1^T7-Cas9^ parental strains was obtained from extracts of ∼ 5x10^6^ PBS-washed cells using the PureLink™ Genomic DNA Mini Kit (Invitrogen). Genomic DNA (1 μg) was submitted for whole genome sequencing by BGI Genomics (100x coverage and approx. 3.5 G reads per sample).

Initial quality control of fastq files was performed using FastQC, followed by adapter trimming and quality filtering with Fastp (0.20.0) ^59^. Processed reads were aligned to the 427_2018 (TriTrypDB v68) genome assembly ^60^ using Bowtie2 (2.3.5) ^61^ with ’--very-sensitive-local’ parameters. The resulting alignments were then processed with SAMtools (1.9) ^62^ for sorting and indexing. Bam files were transformed to bigWig track file or BED format using bamCoverage from DeepTools (3.5) ^71^ with --binSize 5 and --normalizeUsing RPKM. Linear coverage visualization was performed with the svist4get python package ^72^. WT and Δ*ago* bam files were analysed using bamcompare from DeepTools (3.5) ^71^ with --bin 200, smooth 500, --operation log2, --normalizeUsing RPKM –extendReads, --scaleFactorsMethod None, and --outFileFormat bedgraph. The circular fold change visualization of the bedgraph files was performed with the pycirclize (1.6) Python package.

### Machine learning

Differential transcript abundance between 34 °C (control) and 40 °C (heat shock) conditions was obtained from the RNA-seq dataset. Genes with low expression (log2 CPM ≤ 1) were excluded. Protein abundance measurements between 34 °C and 40 °C were obtained from the proteomics dataset. For the analysis, we restricted proteomic measurements to high average expression (log2 intensity > 18). We defined two mutually exclusive categories for each modality (proteome and transcriptome): Upregulated Proteomics (log2 FC > 0.3 and FDR < 0.01), Upregulated Transcriptomics (log2 FC > 0.4 and FDR < 0.01); Downregulated Proteomics (log2FC < −0.55 and FDR < 0.01); Downregulated Transcriptomics (log2 FC < −0.4 and FDR < 0.01) to retain approximately 350 cases in each category. For each gene, we used a recently manually curated version of 5’ UTR and 3’-UTR sequence ^11^. Gene ids were filtered by removing genes with highly similar UTR regions (sequence identity > 40%) using MMseqs2 ^73^. To obtain dense numerical representations of each region, we embedded the 5′-UTR and 3′-UTR sequences using a pretrained HyenaDNA model ^74,75^. from the Hugging Face library LongSafari/hyenadna-medium-450k-seqlen-hf. We trained supervised classifiers to predict the Target label (Upregulated, Downregulated) from sequence embeddings. We repeated this procedure using different subsets of features: all UTR features combined (5′-UTR + 3′-UTR), 3′-UTR-only features, and 5′-UTR-only features. ROC curves from these three feature sets were generated from 3-fold cross-validated probabilities and overlaid for direct comparison. All models were implemented in Python using scikit-learn library.

### Gene Ontology analysis

A GO term enrichment analysis was carried out in R using the ViSEAGO package (version 1.10.0) ^76^. A custom library of terms was prepared by the Custom2GO() function and GO terms associated with each *T. brucei* gene were obtained from TriTypBD (tritrypdb.org) ^60^. GO terms were separated using the create-topGOdata() function (nodeSize =1) and a GO enrichment test was run using the runTest() function (algorithm = “elim”, statistic = “fisher”) prior to collecting the output using the merge_enrich_terms() function. For generation of a heatmap, a semantic similarity analysis was performed using the enrichment analysis data and the build_GO_SS() and compute_SS_distances() functions and then, the heat map was created using the GOterms_heatmap() function (distance = “Wang”, aggreg.method = “centroid” or “ward.D2” and default conditions). The deduced table from the heat map, containing all the relevant information from the GO term analysis and was obtained using the show_table() function.

### Phylogenetic analysis

Trypanosomatid protein sequences were obtained from TryTripDB (tritrypdb.org) ^60^ and a phylogenetic tree was assembled using protein sequences of known members of the HSP60, HSP70, HSP90 and HSP110 families from the *T. brucei* TREU927 reference genome. We searched for orthologs in *Trypanosoma cruzi* Sylvio X10/1-2012 (or Sylvio X10/1 for the HSP70cyt1 and HSP83cyt proteins) and the *Leishmania major* Friedlin genome. Identical or almost identical duplicated genes were not included (one gene ID was selected in each case, see Supplementary Data 1, sheet 3). Multiple sequence alignment including all the protein sequences were carried out using the Clustal Omega web tool (https://www.ebi.ac.uk/Tools/msa/clustalo) ^77^ and the phylogenetic tree was generated usingiTOL (https://itol.embl.de/) ^78^.

### Motif search analysis

Motif enrichment analyses were performed by Multiple Em for Motif Elicitation (MEME) ^79^ comparing the annotated 3’-UTRs in the TREU927 genome ^11^ for cohorts of *T. brucei* genes with a control cohort. The MEME analyses were set up in discriminative mode, to find up to five motifs in the given strand and for a fixed wide of 12 bp. Another similar MEME search was performed to search for palindromes. Only motifs with e-value < 1e-10 were considered. Enriched motifs were used to determine the motif frequency in 5’-UTRs, coding sequences and 3’UTRs of selected cohorts of genes using Find Individual Motif Occurrences (FIMO) analyses ^80^. FIMO analyses were performed by scanning the given strand only using a p-value <1. The results were then manually filtered for p-values as follows: (UUA)_4_ < 1.5 x 10^-5^; A-track < 2.1 x 10^-4^; U-track < 1.5 x 10^-4^; G-track < 3.6 x 10^-4^; C-track < 2.4 x 10^-3^.

### Data processing and visualization

Data processing was performed in a jupyter notebook environment ^81^ using python and R. Omics results were visualized using R packages: ggplot2 (version 4.0.0; https://ggplot2.tidyverse.org), patchwork (version 1.3.2), ggrepel (version 0.9.6), ggpubr (version 0.6.1) scales (version 1.4.0), gghalves (version 0.1.4), or ggVennDiagram (version 1.2.2) ^82^. A karyotype plot for *T. brucei* was created from the Lister427_2018 genome (version 68) obtained from TriTrypDB ^60^ using the circlize package (version 0.4.16). Distance from start and end of polycistons data were from ^31^. Pearson correlation tests were calculated using cor() from base R function, the t-tests were performed using the rstatix package (version 0.7.2). To investigate the relationship between the motif frequencies and gene expression, we employed a Multiple Linear Regression (MLR) model with an interaction term between the frequency of two selected motifs. The dependent variable is the log2 fold change for transcripts or proteins during heat stress, and the independent variables were frequency of motifs. A two-way interaction model was estimated using Ordinary Least Squares (OLS) regression where all parameter tests were conducted using Heteroscedasticity-Consistent Standard Errors (HCSE). The t-statistics and p-values reported in the results are based on the HC3 estimator implemented via the sandwich (version 3.1.1) and lmtest (0.9.40) R packages. Affinity Designer was used to prepare these Figures.

## Data availability

High-throughput sequencing data generated for this study (RNA-seq and whole-genome data) have been deposited in the European Nucleotide Archive, www.ebi.ac.uk/ena under accession code PRJNA1381115 (https://www.ebi.ac.uk/ena/browser/view/PRJNA1381115).

The mass spectrometry proteomics data have been deposited at the PRIDE repository under project code PXD071951 (https://www.ebi.ac.uk/pride/archive/projects/PXD071951).

## Code availability

The code for the proteomics and transcriptomics differential expression analysis and the code for the machine learning analysis have been deposited in GitHub (https://github.com/mtinti/thermoresponse) and archived in Zenodo (mtinti/thermoresponse: v0.1. Zenodo. doi:10.5281/zenodo.18300777).

## ACKNOWLEDGEMENTS

We thank Douglas Lamont and Sam Kosto (FingerPrints Proteomics Facility at the University of Dundee) for assistance with quantitative proteomics.

## AUTHOR CONTRIBUTIONS

G.B.R. Conceptualization, Formal analysis, Investigation, Data curation and visualisation, Writing—original draft, Writing—review and editing; M.T. Data curation, visualisation, and analysis, Writing—review and editing; D.H. Conceptualization, Formal analysis, Funding acquisition, Supervision, Writing—original draft, Writing—review and editing.

